# Synergetic hallmark knockouts immortalize bovine muscle stem cells for cellular agriculture

**DOI:** 10.1101/2025.09.02.673775

**Authors:** Xiaoli Zhang, Benjamin H. Bromberg, Edward B. Gordon, Archana Nagarajan, Andrew J. Stout, Onur Hasturk, Davin Sim, Joy C. Brennan, Nadia D. La, Aneth J. Fernandez, Shlomit David, Eugene C. White, David L. Kaplan

## Abstract

Achieving scalable and sustainable production of cultivated meat hinges on developing robust livestock muscle and fat cell lines that can proliferate and differentiate effectively, while meeting regulatory standards and consumer expectations. In this study, bovine satellite cells (BSCs) were immortalized using CRISPR/Cas9 to knockdown PTEN (phosphatase and tensin homolog), TP53 (cellular tumor antigen) and SMAD4 (SMAD [Sma and Mad proteins] family member 4). The resulting cell line, named “CriBSC2,” exhibited consistent growth, maintained muscle cell characteristics, and successfully differentiated into multinucleated myotubes after more than 150 cell divisions. In contrast, another cell line, “CriBSC1,” achieved immortalization with TP53 knockout alone but lacked differentiation capacity. CriBSC2 were further cultured on gelatin scaffolds to evaluate their anchorage-dependent responses, laying the groundwork for their potential application in tissue engineering for cellular agriculture. Ultimately, CriBSC2 cells demonstrate suitable proliferation and differentiation capabilities crucial for advancing cellular agriculture and future food technologies.

## Introduction

Cells are central to any bioengineering strategy aimed at generating functional tissue for use in regenerative medicine, in vitro model tissue systems, and alternative foods such as cultivated meats. While substantial advances have been realized in regenerative medicine using stem cells and bioengineered versions of these cells for clinical applications, additional progress is needed in fields such as cultivated foods for human health and nutrition.

Cell-cultivated meat is receiving increasing attention to help address growing concerns around industrial agriculture like animal husbandry approaches to food, particularly for protein-rich foods such as meat^1^. Issues associated with conventional animal agriculture include sustainability, animal welfare, antimicrobial resistance, zoonotic diseases, and other factors^2,3^. Cultivated meat, which is produced through large-scale cell culture under controlled conditions, offers the potential for mitigating these externalities while addressing growing global demand for protein-rich foods^2,3^.

Key cell types for the field of cellular agriculture can include embryonic stem cells (ESCs), induced pluripotent stem cells (iPSCs), and primary muscle- or fat-derived stem cells^4,5^. In the latter group, muscle satellite cells (MuSCs) such as bovine satellite cells (BSCs) are particularly attractive due to their robust proliferation capacity and subsequent propensity to differentiate into muscle tissue using known media components^6,7^. Since muscle tissue is crucial for structure and texture in meat, this is a critical focus for the field of cellular agriculture and a promising cell type to pursue^8^. However, a key limitation facing satellite cells such as BSCs for cultivated meat is that they are subject to the Hayflick limit, a point around 50 population doublings after which the cells senesce and lose their ability to divide. To overcome this, immortalized bovine satellite cell lines can be a promising solution, where spontaneous mutations or genetic engineering can overcome senescence, allowing for higher population doubling levels (PDL) while maintaining cellular differentiation capacity, thereby fostering more scalable and economical processes.

Genetic engineering techniques have been previously applied to generate immortalized BSCs (iBSCs), due to difficulties encountered with the spontaneous generation of bovine cell immortalization^9^. These iBSCs were immortalized through random integration of constitutively expressed telomerase reverse transcriptase (TERT) and cyclin-dependent kinase 4 (CDK4), and showed the ability to proliferate and differentiate well past the Hayflick limit (>100 doublings)^10^. However, while these cells are valuable research tools, the use of random integration (through the sleeping beauty system) and the insertion of transgenic nucleotide sequences (such as a puromycin resistance gene) could limit their utility in food. Specifically, these changes would classify the cells as genetically modified (GM) under most regulatory frameworks, thereby posing a significant regulatory barrier. In contrast, targeted gene editing techniques that do not insert foreign DNA (such as cisgenic insertions or non-integrative CRISPER/Cas9 editing) do not trigger GM classification in many regulatory jurisdictions, including the three largest meat-producing countries in the world (the U.S., Brazil, and China)^11^. As such, immortalizing cells via these targeted techniques could present substantially fewer regulatory hurdles. Similarly, consumers often show a preference for foods that are engineered via CRISPR/Cas9 strategies as opposed to conventional genetic modification through transgenes^12^. Together, these facts suggest that CRISPR-edited cells could offer substantial practical benefits over transgenically-immortalized cells.

Here we produced CRISPR immortalized BSCs by specifically downregulating the expression of the PTEN (phosphatase and tensin homolog), TP53 (cellular tumor antigen), SMAD4 (SMAD [Sma and Mad proteins] family member 4) in bovine satellite cells^13–15^, which are essential for numerous cell functions, including cell division and apoptosis^16–18^. As such, it was hypothesized that their downregulation might result in BSC immortalization while retaining the ability of cells to differentiate into multinucleated myotubes. As a proof of concept, the approach and outcomes provide a strategy for using genome editing tools for the immortalization of bovine satellite muscle cells with maintained myogenicity. While regulations for the use of gene-edited cells in cultivated meat remains an open question in many jurisdictions, the recent approval of gene-edited chicken fibroblasts for human consumption in foods suggests that opportunities will exist for the cells and strategies described herein^9,19^.

## Materials and Methods

### 1. Primary bovine satellite cell Isolations, culture, passage, cryopreservation

The methods used for isolation and characterization of the primary bovine satellite cells (BSCs) have been reported in our previous publications (IACU C protocol #G2018-36)^10,20^. Briefly, a 5-week-old Simmental calf at the Tufts Cummings School of Veterinary Medicine, was the source of tissue, and a portion of the semitendinosus muscle was harvested and transported on ice in DMEM+Glutamax (ThermoFisher #10566024, Waltham, MA, USA) with 1% antibiotic/antimycotic (ThermoFisher #1540062) for ∼2 h. The muscle tissue was minced into a fine paste, and then incubated under agitation and regular trituration at 37°C for 45 min with 0.2% collagenase II (Worthington Biochemical #LS004176, Lake wood, NJ, USA) in DMEM+Glutamax (ThermoFisher #10566024, Waltham, MA, USA). Next, the mixtures of the digested tissues were passed through a sterile 18-gauge needle, filtered through 70 µm and 40 µm cell strainers (Sigma #CLS431751-50EA; #CLS431750-50EA, Burlington, MA, USA), and plated at 100,000 cells/cm^2^ in conventional BSC growth medium containing 20% fetal bovine serum (FBS; ThermoFisher #26140079), 1 ng/mL human Fibroblast Growth Factor 2 (FGF-2, PeproTech #100-18B, Rocky Hill, NJ, USA), and 1% Primocin (Invivogen #ant-pm-1, San Diego, CA, USA) in DMEM+Glutamax (ThermoFisher #10566024, Waltham, MA, USA) in an incubator at 37°C with 5% CO_2_. Cell numbers were determined with an NC-200 automated cell counter (Chemometec, Allerod, Denmark). After incubation for 24 h, the cultured media with unattached satellite cells was transferred into new flasks with iMatrix-511 (0.25 μg/cm^2^) (Iwai North America #N892021, San Carlos, CA, USA) to improve cell attachment over three-days of culture. The BSCs were then subjected to conventional culture and passages; usually, cells were passaged at 70% confluency (every 2-3 days) harvesting with 0.25% trypsin-EDTA (ThermoFisher #25200056) and cultured in BSC growth medium with 0.25 μg/cm^2^ of laminin applied with each passage.

For cell cryopreservation, 10% dimethyl sulfoxide (DMSO; Sigma #D2650) in FBS was used. The cryovials containing the cells were placed in a controlled rate freezing apparatus (Corning™ 432005), decreasing the temperature approximately 1°C per minute to –80°C overnight. Then, the frozen cells were transferred to liquid nitrogen. The frozen cells were thawed in a 37°C water bath. Pre-warmed growth media was used to mix with the cryopreserved solution (4:1 vol / vol). The cell pellets were resuspended with fresh growth media after centrifugation at 300 rpm for 5 min and transferred to a new flask for culture. All the procedures were operated in a BSL-2 lab with required PPE.

### 2. CRISPR system and nucleofection

The gRNA sequences (PTEN-gRNA1: GCAGCAATTCACTGTAAAGC; TP53-gRNA1: GTGCGTGTTTGTGCCTGTCC; SMAD4-gRNA1: GAAGGAGAAAAAAGATGAAT; TP53-gRNA2 GTGcAACAGCTCCTGCATGG; SMAD4-gRNA2: GTCAACTCTCCAATGTCCAC) were analyzed via NCBI BLAST to assure that they were identical to the sequence of the *Bos taurus* genome (Taxonomy ID: 9913). Single gRNA vectors of the PTEN-gRNA1 (Addgene 68348); TP53-gRNA1 (Addgene 68349), SMAD4-gRNA1 (Addgene 68350), TP53-gRNA2 (Addgene 68351), SMAD4-gRNA2 (Addgene 68352) and had Cas9 expressed under the CAG promoter. The 5× gRNA vector (Addgene 68357) included the five single gRNA targets of the TP53, SMAD4 and PTEN but without Cas9. All five single gRNAs and 5× gRNA vectors were gifts from Jannik Elverløv-Jakobsen (Department of Biomedicine, Aarhus University). The vector for expressing Cas9 without gRNAs was a gift from Timo Otonkoski (Helsingin Yliopisto, Finland) purchased from Addgene (Addgene 78311)^21^. These vectors were used for BSC nucleofection. Plasmids for six engineered cell lines (TP53-gRNA1, TP53-gRNA2, SMAD4-gRNA1, SMAD4-gRNA2, PTEN-gRNA1 and PTEN/TP53/SMAD4-5×gRNAs cell lines) mentioned above were prepared with high concentrations (1 ug/ ul) and then subjected to nucleofection using Lonza 4D-Nucleofector™ system (Lonza, Australia). The parameters for the nucleofection were as follows: program EY-100, P5 solution, 18-strip cuvettes and 50,000 cells for each reaction. Different from the single gRNA vectors, the 5× gRNA vector was transfected together with the vector for Cas9 expression (Addgene 78311), which was designated as PTEN/TP53/SMAD4-5× gRNAs cell line. To verify the indels caused by the CRISPR cuts in the six CRISPR engineered cell lines, primers listed in supplementary Table 1 were used for PCR clones and NGS analysis (Table S1).

### 3. Establishing immortalized BSCs through long-term cell passage

After the nucleofection, the cells were cultured with BSC growth media as before. Before cell growth stabilized, the engineered cells experienced several survival crises, which was followed by long-term culture^10^. The bovine satellite cells in each of the six engineered groups (TP53- gRNA1, TP53- gRNA2, SMAD4- gRNA1, SMAD4- gRNA2, PTEN- gRNA1 and 5× gRNA-Cas9 group) were cultured under the same conditions as the primary BSCs with plating densities of 4,000 cells/cm^2^ in T75 flasks. The reagent iMatrix laminin-511 (Iwai North America #N892021, San Carlos, CA, USA), was used as substrate at every passage (0.25 μg / cm^2^). When cell confluency reached 70 to 80%, the cells were harvested with 0.25% trypsin-EDTA (ThermoFisher #25200056). Cell numbers were counted with an NC-200 automated cell counter (Chemometec, Allerod, Denmark) and recorded for long-term culture curves.

In the CRISPR engineered groups, cells from TP53- gRNA1, SMAD4- gRNA1, SMAD4- gRNA2 and PTEN- gRNA1 groups ceased growth at passage 37 (P37) with 91 doublings, P30 with 80 doublings, P32 with 76 doublings, and P29 with only 51 doublings, respectively. The engineered cells from TP53- gRNA1 reached 150 doublings at P52, while the engineered cells from 5× gRNA-Cas9 achieved 163 doublings at P50. Hence, the two engineered groups, CriBSC1 (TP53- gRNA1) and CriBSC2 (5× gRNA-Cas9) were designated as successfully immortalized cells (over 100 doublings).

### 4. Immunostaining of the immortalized CriBSC1 and CriBSC2

To analyze proliferative cell phenotype, cells from the immortalized CriBSC1 (P30 – 39, 100 - 120 cumulative doublings), CriBSC2 (P30 – 35, 100 - 120 cumulative doublings), and non-engineered primary (P2 - P5) BSCs, were cultured with conventional growth media in 48 well plates to 70% confluency. Then, the plates were fixed and subjected to immunostaining with primary antibody Pax7 (Rabbit, ThermoFisher #PA5-68506, Dilution 1:500), Phalloidin 488 (Invitrogen™ A12379, Dilution 1: 1,000), MyoD (Mouse, ThermoFisher #MA5-12902, Dilution 1:100), and Myogenin (Mouse, Santa Cruz #sc-52903, Dilution 1:250). To analyze cell differentiation, cells from each engineered group were cultured in triplicate in 48 well plates to 70% confluency. After that, the cells were cultured for about 3 – 5 days with differentiation medium media including 2% fetal bovine serum (FBS; ThermoFisher #26140079), 1 ng/mL human Fibroblast Growth Factor 2 (FGF-2, PeproTech #100-18B, Rocky Hill, NJ, USA), and 1% Antibiotic-Antimycotic (ThermoFisher #1540062) in DMEM+Glutamax (ThermoFisher #10566024, Waltham, MA, USA) in an incubator at 37°C with 5% CO_2_. Next, the cells were fixed and subjected to immunostaining with primary antibody Myogenin (Rabbit, ThermoFisher #PA5-116750, Dilution 1:20) and MF20 (Mouse, R&D Systems MAB4470SP, dilution to 4ug/mL final concentration). Secondary antibodies used for both proliferation and differentiation were Goat-anti-Mouse 488 (ThermoFisher #A11001, Dilution 1: 1,000) and Goat-anti-Rabbit 594 (ThermoFisher #A11072, Dilution 1:500). A blue-fluorescent DNA stain, DAPI (1 mg/ml, Thermo Scientific™ 62248, Dilution 1: 1,000) was used to stain the cell nucleus. Each experimental group had three replicates.

The immunostaining procedures were 1) Fixation & Permeabilization: The cells were fixed for immunostaining at room temperature with 4% paraformaldehyde (PFA, ThermoFisher #AAJ61899AK) for at least 30 minutes after removing the culture media. The samples were then rinsed three times with DPBS (ThermoFisher #14190250), permeabilized for 15 min with 0.5% Triton-X (Sigma # T8787), rinsed three times with PBST with 0.1% Tween-20 (Sigma #P1379) in PBS and blocked for 45 minutes with blocking buffer including 5% goat serum (ThermoFisher #16210064) and 0.05% sodium azide (Sigma #S2002) in DPBS. 2) Primary stain: The blocking buffer from step 1 was aspirated and the samples were washed three times with PBST. The diluted primary antibody solutions were incubated at 4°C overnight. 3) Secondary stain: The primary antibody solutions were removed, and the samples were rinsed with PBST three times. The cells were soaked in Wash Buffer for 15 minutes. A dilute secondary antibody was added to the wash buffer and incubated in the dark at room temperature for 60 minutes. 4) Imaging: The secondary antibodies were removed, and the samples were rinsed twice with PBS and then soaked in PBS for 5 minutes prior to imaging. DAPI was added while incubating with the secondary antibodies. All the experiments were performed in a BSL-2 lab with appropriate PPE and in a fume hood. Images were analyzed under BZ-X810 fluorescent microscope (Osaka, Japan).

### 5. Gene expression in immortalized CriBSC1 and CriBSC2 (qPCR)

To assess gene expression patterns in the immortalized CriBSC1 (P30 – 39, 100 - 120 cumulative doublings) and CriBSC2 (P30 – 35, 100 - 120 cumulative doublings) non-engineered primary (P2 - P5) BSCs, primers for the housekeeping gene 18S (ThermoFisher #Hs03003631) were used as inner reference. Primers for Pax3 (ThermoFisher #Bt04303789), MyoD1 (ThermoFisher #Bt03244740), Myogenin (ThermoFisher #Bt03258929), and Myosin Heavy Chain (ThermoFisher #Bt03273061) were used to investigate proliferation and differentiation of the immortalized BSCs. Primers for TP53 (ThermoFisher #Bt03223218_g1), SMAD4 (ThermoFisher #Bt03243052_m1), and PTEN (ThermoFisher #Ss03820741_s1) were used for detecting expression levels of these genes in the engineered cells. To perform RT-PCR, RNA was extracted at 70% cell confluency or after differentiation using the RNEasy Mini kit ((Qiagen #74104, Hilden, Germany) and cDNA was synthesized using the iScript cDNA synthesis kit (Bio-Rad #1708890) according to the manufacturer’s instructions. qPCR was performed using 2 μL of cDNA and 1 μL of each primer (mentioned above) in a 20 μL total volume per reaction (TaqMan™ Fast Universal PCR Master Mix (2X), no AmpErase™ UNG, cat. 4366072). Reactions were performed on a CFX96 Real Time System thermocycler (Bio-Rad, Hercules, CA, USA), and the run information included initiation (10 minutes at 95°C) with 49 repeats (step 1 at 95°C for 15 seconds and step 2 at 60°C for 60 seconds). Relative quantification of the qPCR results was analyzed as 2^-ΔΔct^.

### 6. RNA-Seq library preparation and sequencing

RNA was extracted and isolated from three biological replicates of proliferating (70% confluency) primary BSCs (passage 4) and CriBSC2 at passages 16, 30, and 47 (50,100, and 150 cumulative doublings respectively) using a RNEasy Mini kit (Qiagen #74104, Hilden, Germany) according to manufacturer’s directions. RNA samples were submitted to Azenta Genewiz for transcriptome library construction and Illumina sequencing. Library preparation was conducted using Genewiz’s Standard RNA-Seq Service targeting mRNA with Poly(A) selection for RNA samples with an RNA Integrity Number (RIN) score of 6 or better. Sequencing was performed on Illumina NovaSeq 6000 S4 with paired-end, 150 base pair (bp) read.

### 7. Read alignment and quantification

Scripts for read alignment and quantification of sequencing data were adapted from the open access materials provided by the teaching team at the Harvard Chan Bioinformatics Core^22^. Unless noted otherwise, for all software used, all parameters not mentioned specifically were set to default values. Quality control (QC) was performed for raw reads using FastQC^23^ (v0.12.1). Over-represented sequences were searched against the NIH nucleotide collection (nt) using BLASTN and TBLASTN^24^ for alignment to the *Bos taurus* genome to identify adapter sequences or contamination. For each sequencing library, read alignment QC was performed using STAR^25^ with a STAR genome index (parameters: -- runThreadN 12 --runMode genomeGenerate --sjdbOverhang 149). Both STAR alignment and STAR genome index generation used the NCBI RefSeq assembly of the *Bos taurus* reference genome bosTau9 (ARS-UCD2.0). Qualimap^26,27^ was subsequently used to analyze the read alignment QC, as well as determine the genomic origin of reads and transcript coverage. Informed by biases found in the alignment QC analysis, pseudo-alignment was performed using Salmon^28^ (v1.10.1, parameters: quant -p 24 -l A --seqBias --posBias --validateMappings --writeUnmappedNames) to compute psuedocounts for transcript quantification. The selective alignment mapping strategy^29^ for Salmon was used for transcript quantification with a decoy-aware transcriptome index (parameters: index -p 12 -k 31) to mitigate spurious mapping of reads during quantification. An HTML report summarizing the raw read QC, read alignment QC, and transcript quantification results was generated using MultiQC^30^ (v1.18) for multi-sample comparison. This work was done on the Tufts High-Performance Computing Research Cluster using the environment manager miniconda^31^ (v23.10.0).

### 8. Transcriptome analysis

Scripts for transcriptome analysis of counts data were adapted from the open access materials provided by the teaching team at the Harvard Chan Bioinformatics Core^22^. Using the DESeq2^32^ package (v1.42.0), raw Salmon pseudocounts were normalized by the median of ratios method and log2 transformed through a Regularized Log Transformation (rlog) for visualization and clustering of read count data. Sample-level QC was subsequently performed on the log2 transformed normalized counts. Using DESeq2^32^, principal component analysis was performed using singular value decomposition on the 1000 most variable genes to examine the co-variances between samples. To further explore the co-variances between samples, unsupervised hierarchical clustering was performed using the pheatmap^33^ CRAN package (v1.0.12). Complete-linkage analysis with a Euclidean distance measure was used for clustering.

Dispersion estimation using DESeq2 was calculated for normalized count data and estimates were shrunken towards a parametric trendline before performing Wald statistics. Differential gene expression (DGE) was performed using a Wald test with a Benjamini-Hochberg False-Discovery Rate^34^ (BH FDR) significance level of 0.05 and a log2 fold change (log2FC) threshold of 1. We tested for differences due to cell type by assessing the differential expression of genes in CriBSC2 at each time point (passage 16, 30, or 47) relative to primary BSCs. Log2FC results from DGE were shrunken toward zero for genes with low counts and/or high dispersion values using the ashr method^35^. Additionally, DGE results were explored using volcano plots, created using the EnhancedVolcano^36^ Bioconductor package (v1.13.2). GSEA was performed on a log2 fold change sorted list of all differentially expressed genes (DEGs). GSEA was run against the Gene Ontology (GO) gene set for *Bos taurus* using the clusterProfiler^37,38^ (v4.10.0) Bioconductor package with a seed of 12345. We identified significantly enriched pathways using a BH FDR^34^ significance level of 0.05. Analysis was performed using R Statistical Software (v4.3.2; R Core Team 2023).

### 9. Immortalized BSCs cultured on gelatin scaffolds

Biomaterial films (surface patterned and un-patterned gelatin) were prepared as previously described with slight modifications^39^. Briefly, 8 mm diameter round polydimethylsiloxane (PDMS) disks with 1 - 2 mm thickness was prepared by casting onto a reflective diffraction grating with individual groove widths of 1,600 nm and a blaze angle of 28.68° (Edmund Optics, Inc, #55–259, Barrington, NJ). For non-patterned PDMS disks, the cast PDMS substrates were punched into round disks of 8 mm diameter, followed by washing in 70 % ethanol and then rinsing in DI water. These substrates were then used for casting the gelatin solutions to generate patterned films. The non-patterned PDMS substrates were prepared by casting PDMS onto petri dishes and punching round disks of 8 mm diameter. For each substrate, 200 μL solution was cast onto the patterned and nonpatterned PDMS substrates.

Gelatin films^39^: A gelatin solution (5 wt%) was prepared by dissolving gelatin powder (bovine skin, Sigma, USA) in water at 65°C for 2 h, cooling to near room temperature and mixing in 1 wt% microbial transglutaminase (Ajinomoto Ti, Japan). The solution was incubated at 37°C for 1 h to initiate cross-linking^40^. This solution was cast onto the PDMS substrates and dried at 60°C. The films were sterilized by exposure to UV light for 1 h^39^. Following UV sterilization, films were soaked in media for 24 hours followed by seeding at 5,000 cells/cm^2^.

Gelatin salt leach porous sponges were prepared using 20% gelatin powder (from bovine skin, Sigma, USA), 30 U microbial transglutaminase (mTG) (Ajinomoto Ti, Japan) per gram of gelatin, and 600-710 nm NaCl (CAS No. 7647-14-5, Sigma, USA) at a ratio of 2 g salt per 1mL of gelatin/mTG solution. This was mixed in thoroughly, and the solution was added to PDMS cylinder molds of 6 mm diameter by 6mm height, then placed at 37°C overnight for cross-linking followed by leaching in DI water for 3 days with 2-3 changes per day. Scaffolds were autoclaved in water at 121°C for 15 minutes^39^. Following sterilization, scaffolds were soaked in media for 24 hours prior to seeding. The seeding cell density was at 5,000 cells/sponge.

The metabolic activity of the bovine satellite cells cultured on the films and in the sponges was determined at days 1, 7, 14, 21, and 28 via an AlamarBlue HS viability assay (ThermoFisher, A50101) according to the manufacturer’s instructions. Briefly, after rinsing with DPBS, the cells cultured on films were incubated in 400 μL and sponges were incubated in 800 uL of 10 % AlamarBlue reagent (in growth media for 4 hours at 37°C with 5% CO_2_. Following incubation, 100 μL aliquots were transferred into 96-well plates and measured with Excitation/Emission at 560/590 nm using a microplate reader (BioTek Synergy, USA). Assays were performed in triplicate.

Immunofluorescent staining of bovine satellite cells (BSCs) was used to visualize growth and morphology on the films and in the sponges. BSCs were seeded on the films and sponges with a density of 5,000 cells/film or 5,000 cells/sponge in 48 ultra-low attachment well plates (Corning™ 3473, USA). To induce myogenic differentiation, BSCs were cultured for 14 days in growth medium and then differentiated in differentiation medium for 14 days, as described in our previous work^39^. The fixation, permeabilization and blocking of BSCs on films and sponges were performed using 4 % paraformaldehyde for 45 minutes, permeabilization using 0.5 % Triton X‐100 in DPBS for 45 minutes and blocked with 3% bovine serum albumin for 1 h. The samples were incubated with 1:400 AlexaFluor™ 488 phalloidin (Invitrogen) and 1ug/mL DAPI. Confocal laser scanning microscopy (CLSM) imaging was performed on a TCS SP8 microscope from Leica Microsystems (Wetzlar, Germany).

### 10. Statistical Analysis for In Vitro Culture

GraphPad Prism 9.0 software (San Diego, CA, USA) was used for statistical analysis. Doubling time and gene expression were analyzed via one-way ANOVA. Multiple comparisons were performed with Tukey’s HSD post-hoc test with comparisons between all samples. Gene expression was analyzed via unpaired t tests. P values <0.05 were treated as significant. All error bars represent ± standard deviation.

## Results

### 1. Establishment of immortalized bovine satellite cell lines by CRISPR engineering

After isolation from a five-week-old Simmental calf, primary bovine satellite cells (BSCs) (passage 2 - 5) were immortalized by incorporating CRISPR plasmids into the cells (Fig.1a). These plasmids targeted phosphatase and tensin homolog (PTEN), cellular tumor antigen (TP53) and SMAD (Sma and Mad proteins) family member 4 (SMAD4) with five different gRNAs (Fig. 1b). For PTEN inactivation, one gRNA was selected to target the protein tyrosine phosphatase-like catalytic domain. For TP53 inactivation, two gRNAs were selected to target the seventh exon (gRNA2) to destroy the DNA binding site and the eighth exon (gRNA1) to destroy the DNA interaction site. For SMAD4 inactivation, two gRNAs were selected to target the first exon (gRNA1) to destroy the interaction with TSC22 Domain Family Member 1 (TSC22D1), which regulates the transcription of multiple genes, and the eighth exon (gRNA2) to destroy the trimer interface, resulting in loss of function in polypeptide binding. Consequently, five different CRISPR engineered cell lines with single gRNA (PTEN-gRNA1, TP53-gRNA1, SMAD4-gRNA1, TP53-gRNA2, SMAD4-gRNA2) were generated. Additionally, the BSCs engineered with the five single gRNAs all in one plasmid, were termed the PTEN/TP53/SMAD4-5×gRNAs cell lines. The genome of the six CRISPR engineered cell lines were extracted after nucleofection for 24-72 h and were screened for mutations through NGS. Mutation analysis confirmed successful mutations, yielding heterogeneous populations of engineered cells (Fig. 1).

**Fig. 1.**
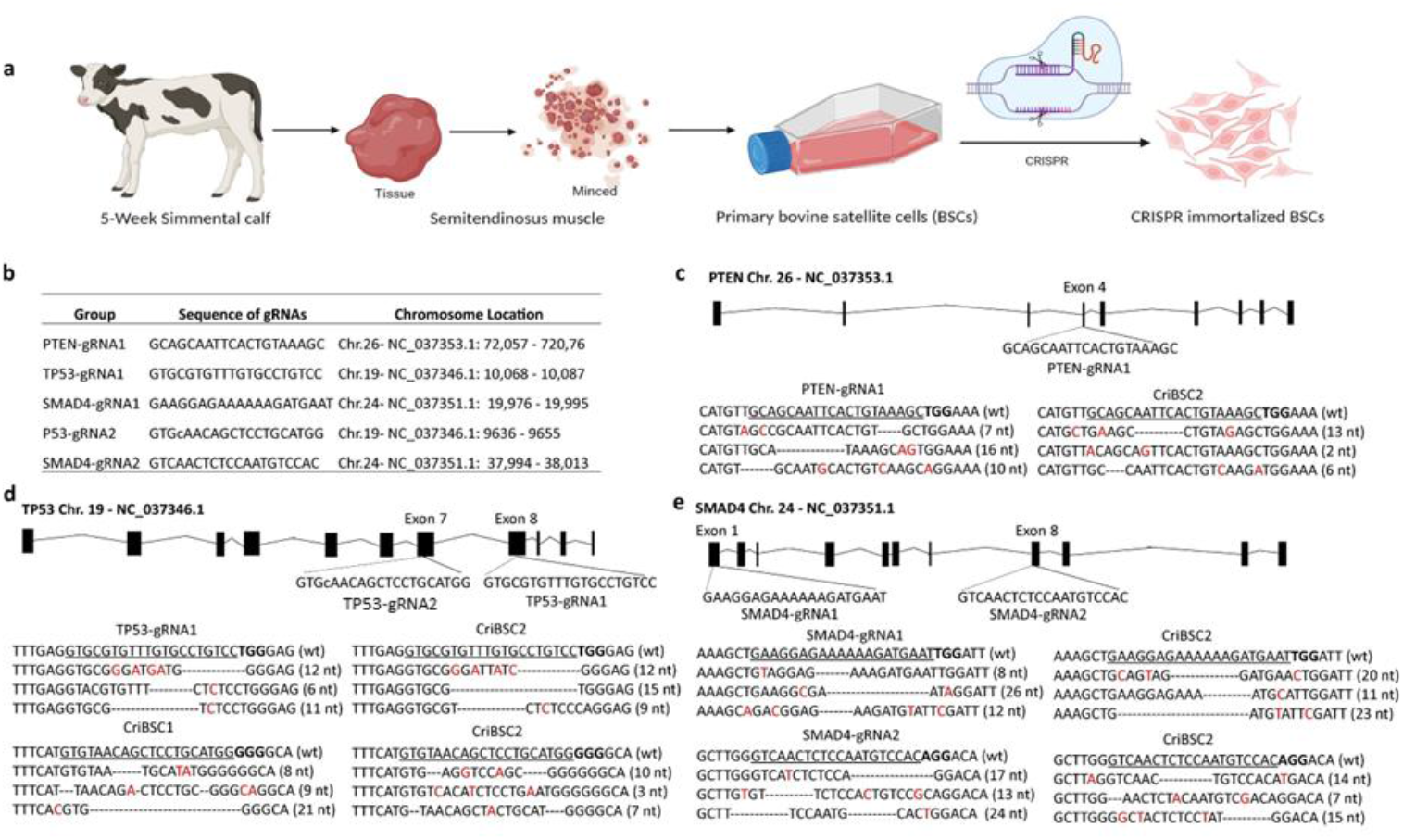
Immortalization of bovine satellite cells by CRISPR engineering: a, Strategy for developing the CRISPR immortalized bovine satellite cell lines. b, Sequence of the five single gRNAs and their chromosome targeting locations. c, Representative mutations of PTEN in both the PTEN-gRNA1 and PTEN/TP53/SMAD4-5×gRNAs cell lines. d, Representative mutations of TP53 in the TP53-gRNA1, TP53-gRNA2 and PTEN/TP53/SMAD4-5×gRNAs cell lines. e, Representative mutations of SMAD4 in the SMAD4-gRNA1, SMAD4-gRNA2 and PTEN/TP53/SMAD4-5×gRNAs.

The CRISPR engineered cell lines initially experienced several survival crises before their growth gradually stabilized. Through long-term culture, the successfully immortalized cell lines, CriBSC1 (TP53-gRNA2) and CriBSC2 (PTEN/TP53/SMAD4-5×gRNAs), passed 100 total population doublings (TPD) at passages 30 and 29, respectively (Fig. 2c). The morphology of the cells in CriBSC1 and CriBSC2 maintained a patch-like morphology as with the primary BSCs (Fig. 2a). In contrast, the unsuccessfully immortalized cell lines (PTEN-gRNA1, SMAD4-gRNA1, TP53-gRNA2, and SMAD4-gRNA2) exhibited spindle-like elongation and network formation (Fig. S2a). The average cell diameters in the CRISPR engineered cell lines were 15 – 16 µm, showing no significant differences compared to that of the primary BSCs (Fig.2b). The PTEN-gRNA1, TP53-gRNA1, SMAD4-gRNA1, and SMAD4-gRNA2 cell lines discontinued proliferation at 51, 91, 79, and 76 TPD, respectively (Fig. S2b). The TP53-gRNA1 cell line with mutation at the DNA interaction region stopped proliferating at passage 38 with 91 TPD, whereas the TP53-gRNA2 cell line, with mutation at the DNA binding region, achieved 150 TPD (Fig. 2c). Both the SMAD4-gRNA1 and the SMAD4-gRNA2 cell lines ceased proliferation after approximately 80 TPD (Fig. Sb). The PTEN-gRNA1 cell line had only 50 TPD as with the primary BSCs. The average doubling time for CriBSC2 (36.6 h) was faster than the primary BSC control (53 h), which is likely due in part to the gradual senescence and associated slow-down of primary BSC growth over multiple passages (Fig. 2e). The average doubling time in other CRISPR engineered groups showed no significant difference compared to the primary BSCs (Fig. 2d and 2f). The CriBSC1 and CriBSC2 had doubling times of about 24 - 48 h as the fastest-growing cell lines (Fig. 2g and 2h).

**Fig. 2.**
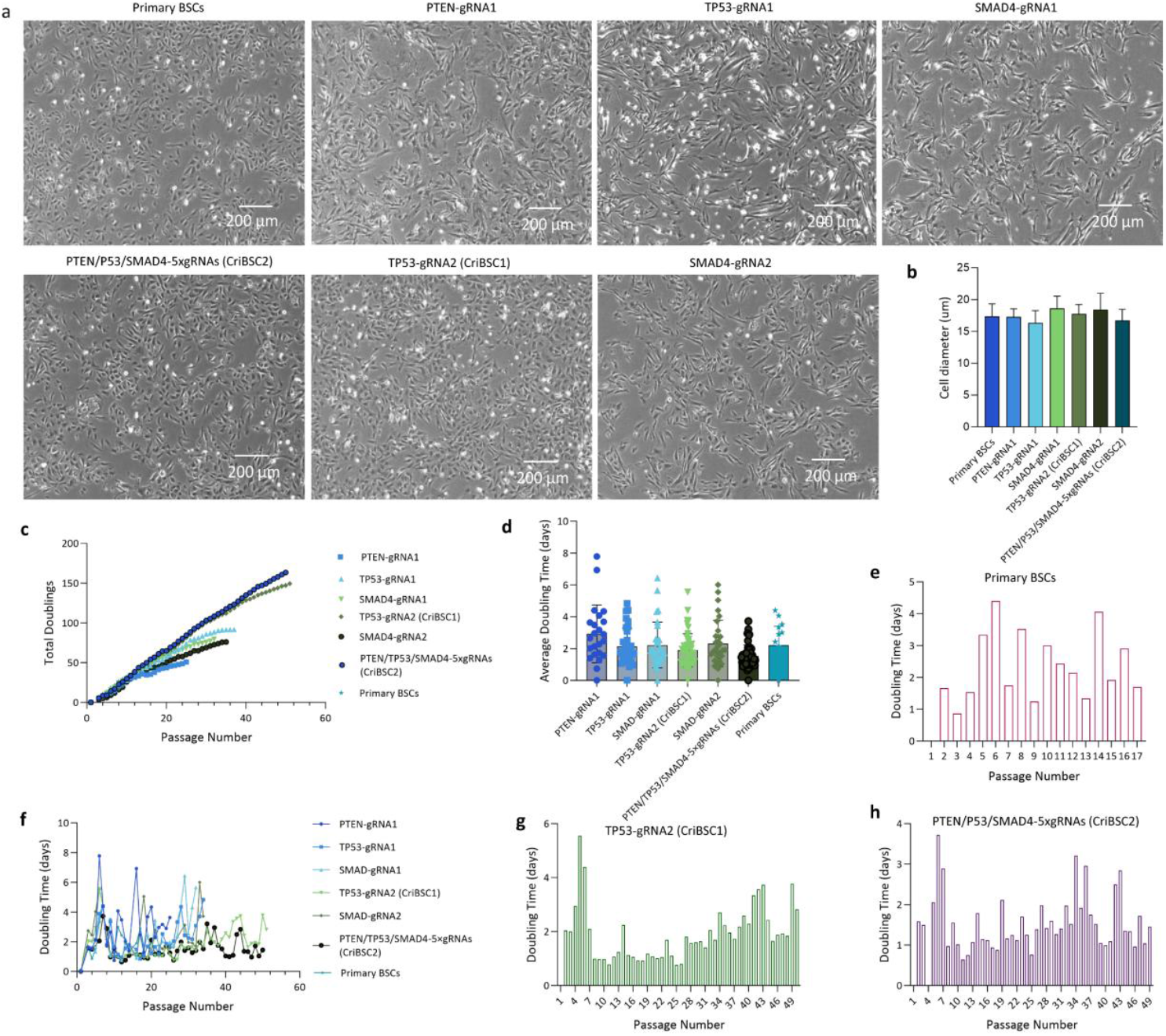
Establishment of CRISPR immortalized bovine satellite cell lines. a, Morphology of the bovine satellite cells for the PTEN-gRNA1, TP53-gRNA1, SMAD4-gRNA1, TP53-gRNA2 (CriBSC1), SMAD4-gRNA2, and PTEN/TP53/SMAD4-5×gRNAs (CriBSC2) compared to primary BSCs. b, cell diameter size for each cell line. c, Total cell population doubling numbers. d, Average doubling time (days). e, Doubling time in the primary BSC control cell line. f, Overall doubling time (days). g, Doubling time in the TP53-gRNA2 cell line (CriBSC1). h. Doubling time in the PTEN/TP53/SMAD4-5×gRNAs cell line (CriBSC2).

### 2. Proliferation and differentiation of immortalized bovine satellite cell lines

After immortalization, cells from CriBSC1 and CriBSC2 were subjected to immunostaining and qPCR to investigate their proliferation and differentiation capabilities. Immunostaining revealed that the expression levels of early satellite cell makers, the paired box homeotic gene 7 (Pax7) and the myoblast determination protein 1 (MyoD), which are associated with proliferation, had decreased. Conversely, the differentiation markers Myogenin and Myosin heavy chain (MyHC) were almost undetectable in CriBSC1 (Fig. 3a and 3b). These findings indicated that CriBSC1 retains its proliferative ability but loses its differentiation potential. In CriBSC2, the expression levels of MyoD, MyoG, Myogenin, and MyHC were suppressed compared to primary BSCs, but still higher than in CriBSC1. This suggests that CriBSC2 retains the capability for both proliferation and differentiation. (Fig. 3a and 3b).

**Fig. 3.**
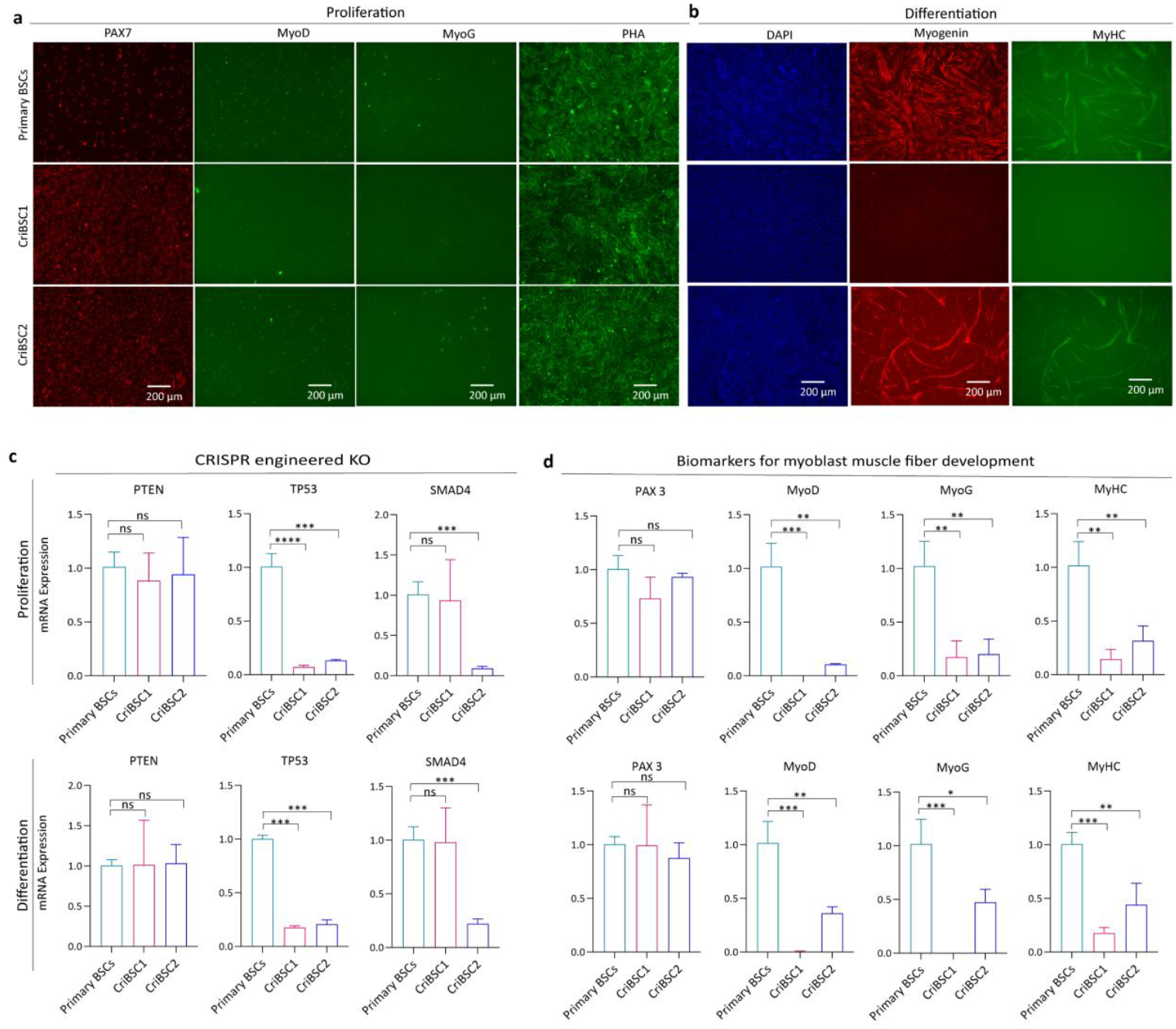
Proliferation and differentiation of immortalized satellite cells. a, Immunostaining of proliferative and b, differentiated cells for the biomarkers PAX7, MyoD, MyoG and MyHC for myogenic progression in CriBSC1 and CriBSC2 compared to the primary BSCs. Cells are also stained with with F-actin (a) and DAPI (b) to indicate cytoskeleton and nuclei, respectively. Scale bars are 200 µm. b, Gene expression levels of c, the CRISPR engineered knockout genes (TP53, PTEN and SMAD4), and d, biomarkers (PAX3, MyoD, MyoG, and MyHC) in CriBSC1 and CriBSC2 compared to the primary BSCs. Error bars indicated s.d. n=3. * p≤0.05; ** p≤0.01, *** p≤0.001.

Gene expression levels of the proliferative and differentiative cells of CriBSC1 and CriBSC2 were investigated using qPCR, with primary BSCs as a control (Fig. 3c - 3e). Analyzing the engineering targets, the expression level of TP53 was significantly decreased in both CriBSC1 and CriBSC2 compared to the primary BSCs (Fig. 3c). Additionally, the expression level of SMAD4 was significantly decreased in CriBSC2, indicating the effectiveness of the cocktail knockouts in this cell line (Fig. 3c). Interestingly, the expression level of PTEN remained unchanged in both CriBSC1 and CriBSC2 (Fig. 3c). Here, while PTEN mutations did not result in reduced RNA transcription, it is possible either that the mutations present did not cause a reduction in PTEN function (no reduction in transcripts or in protein function), or that they did cause a reduction manifested at the protein level rather than the transcription level (therefore undetectable from RNA transcription data). Because of this, while the degree of transcription of PTEN remains unchanged in CriBSC2, the degree of function of PTEN remains unknown (and could be an interesting area for future work). Ultimately, though, these results suggest that TP53 knockdown successfully overcomes cellular senescence but may compromise differentiation capacity, as indicated by the immunostaining data. At the same time, simultaneous knockdown of TP53 along with at least SMAD4 can generate immortalized and myogenic cells. This hypothesis is further supported by the higher expression levels of the MyoD and MyoG in CriBSC2 compared to CriBSC1 (Fig. 3d). Similarly, the elevated expression level of the myosin heavy chain (MyHC) in CriBSC2 compared with CriBSC1 reinforces this hypothesis (Fig. 3d).

### 3. Transcriptomic Analysis of CriBSC2

To gain insights into the transcriptomic changes occurring in BSCs following CRISPR engineering, we conducted bulk RNA-Seq analysis on proliferating CriBSC2 at passages 16, 30, and 47 (corresponding to 50, 100, and 150 cumulative doublings, respectively), with primary BSCs at passage 4 serving as comparisons. Subsequent exploratory read alignment by STAR to the *Bos taurus* reference genome resulted in >93% mapping for reads in all samples. This was accounted for during Salmon transcript quantification, which resulted in >77% mapping for reads in all samples. Sample-level QC by principal component analysis (PCA) and hierarchal clustering of log2 transformed normalized salmon pseudo counts demonstrated expected clustering of biological replicates and indicated no significant outliers and/or sample contamination. PCA revealed that 82% of the variance between the clusters was accounted for by PC1, which appeared to capture the differences induced by the CRISPR knockout treatment. Meanwhile, PC2 explained 12% of the variance, reflecting the variation related to the time that had elapsed since the CRISPR knockout treatment, distinguishing CriBSC2 at passage 16 from those at passages 30 and 47. Hierarchal clustering also outlined the same temporal variation seen in PC2 (see Fig. 4a and 4c). Dispersion estimation using DESeq2 indicated that our data was a good fit for the negative binomial model that is required by DESeq2 analysis (see Fig. 4b).

**Fig. 4.**
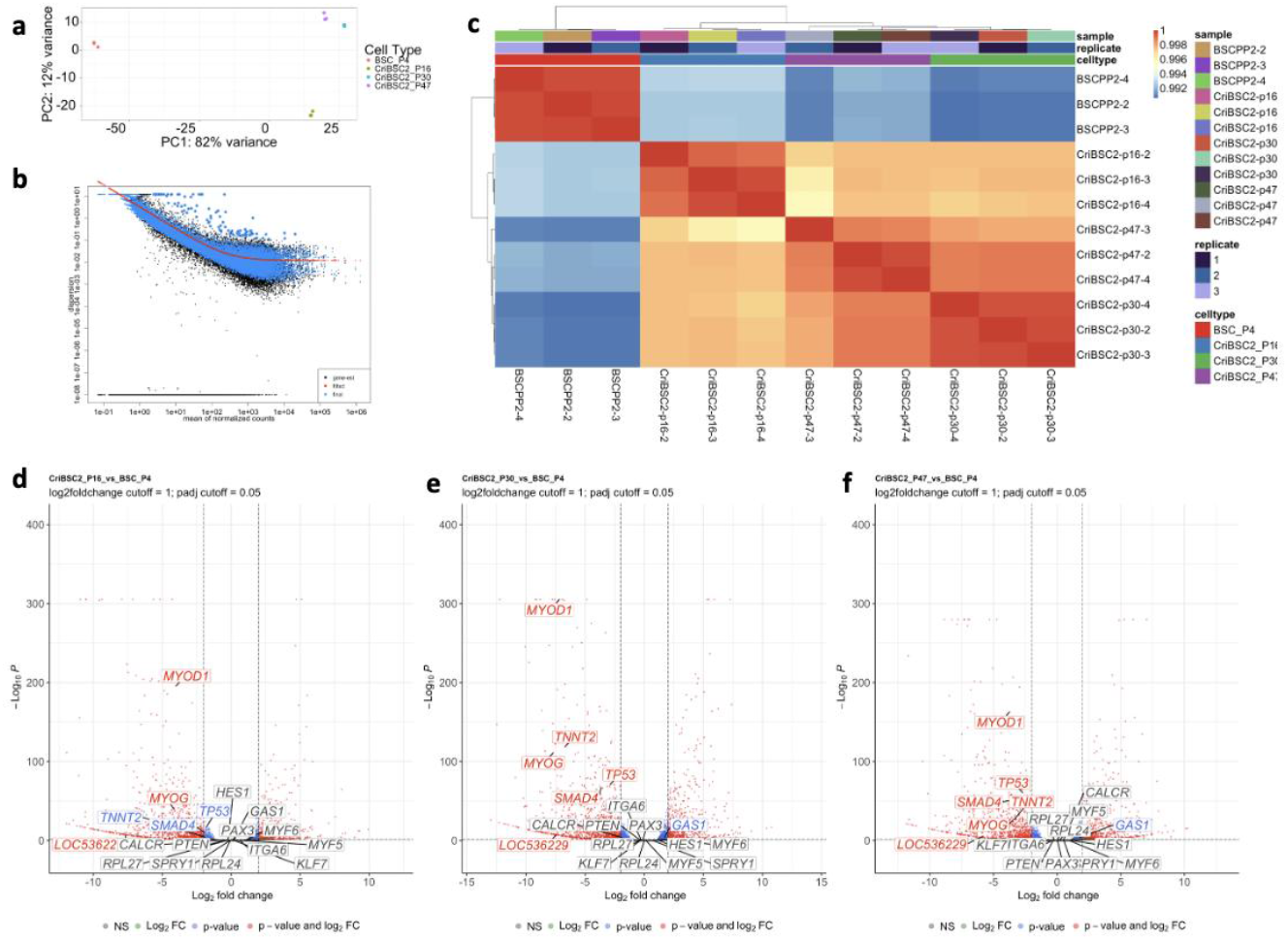
Differential expression analysis of CriBSC2 and primary BSCs by RNA-Seq. a, Principal component analysis by cell type of log2 transformed normalized salmon pseudocounts of the 1000 most variable genes. **b**, DESeq2 dispersion estimates of log2 normalized salmon pseudocounts. **c**, Unsupervised hierarchical clustering of differential gene expression results. Results are clustered using the ward. D2 algorithm based on a Euclidean distance measure. Z-score statistics were calculated on a gene-by-gene basis after clustering. **d**, Volcano plot of DGE results based on a Benjamini-Hochberg False-Discovery Rate (BH FDR) significance level of 0.005 and a log2FC cut-off of 0.585. P-values correspond to BH FDR adjusted p-values. Genes of interest are labeled and colored according to significance.

Differential gene expression was performed using DESeq2 with a Benjamini-Hochberg False-Discovery Rate (BH FDR) significance level of 0.05 and a log2 fold change threshold of 1 (see Table S2). Differential gene expression results were explored and revealed a notable downregulation of many of the canonical biomarkers associated with early-stage MuSC activation and myogenic commitment (MYOD1) and late-stage MuSC activation / myoblast differentiation (MYOG and TNNT2) in CriBSC2 compared to primary BSCs for all CriBSC2 passage timepoints (passage 16, 30, or 47). Notably, early-stage MuSC activation and myogenic commitment markers MYF5, RPL24, and RPL27 and late-stage MuSC activation / myoblast differentiation MYF6/MRF4 did not exhibit significant upregulation or downregulation. Many biomarkers associated with MuSCs in a quiescent or close-to-quiescent state (PAX3, SPRY1, HES1, GAS1, CALCR, KLF7) did not exhibit significant upregulation or downregulation. Interestingly, the uncharacterized annotation for paired box protein Pax-7 (LOC536229), a MuSC marker for quiescence that is downregulated upon activation, was found to be downregulated in all CriBSC2 passage timepoints. Moreover, our differential expression results indicated that transcripts of TP53 and SMAD4 were downregulated, whereas PTEN did not exhibit significant upregulation or downregulation according to our BH FDR and log2 fold change thresholds (see Fig. 4d-f and Table S2). Gene set enrichment analysis (GSEA) was performed against the Gene Ontology (GO) gene set for *Bos taurus* to identify differentially expressed biological pathways in CriBSC2 compared to primary BSCs. GSEA demonstrated an enrichment and overall, up-regulation for pathways related to cell cycle and cell replication mechanisms in CriBSC2 compared to primary BSCs for all CriBSC2 passage timepoints at a BH FDR significance level of 0.05 (see Table S2).

### 4. Long-term culture on 3D gelatin scaffold of CriBSC2

Culturing CriBSC2 on gelatin films and scaffolds showed better coverage than primary BSCs after 28 days (Fig. 5a and 5b). F-actin staining showed large portions of surface coverage on both nanopatterened films and salt leach sponges (Fig. 5a and 5b). Metabolic activity, assessed using AlamarBlue and cell proliferation, measured by Cyquant analysis, showed significant improvements in the growth of CriBSC2on 3D sponges. The metabolic activity of CriBSC2 was nearly twice that of primary cells, and the double-stranded DNA contant was eight times greater (Fig. 5c and 5d). Furthermore, over the 28-day culture period, Cyquant analysis indicated sustained growth of CriBSC2 on both the salt-leach gelatin sponges and nanopatterned gelatin films (Fig. 5d).

**Fig.5.**
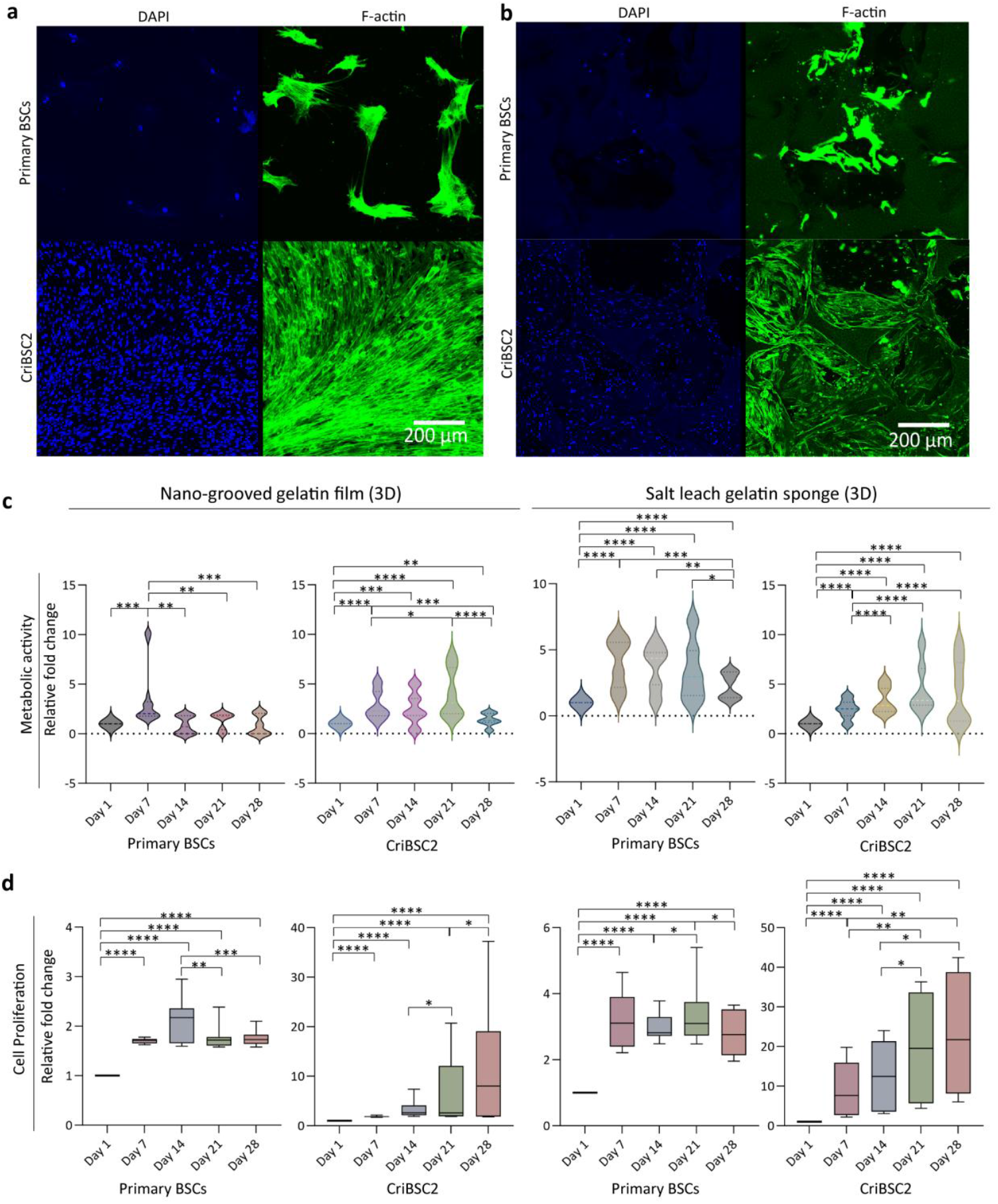
Culture of CriBSC2 and primary BSCs on 3D gelatin scaffolds. a and b, Immunostaining of nuclei (DAPI) and actin (F-actin) of CriBSC2 and primary BSCs on gelatin nano-grooved films (a) and salt leach sponges (b). Scale bars are 200 um. c and d, the metabolic activity (Alamar Blue) and the cell proliferation (Cyquant) of the CriBSC2 and primary BSCs cultured on nano-grooved films and salt leach sponges through day 1 to day 28. Error bars indicated s.d. n=3. * p <0.05; ** p<0.01, *** p<0.001, **** p<0.0001.

## Discussion

Establishing immortalized cell lines that maintain their differentiation capacity is essential for advancing muscle and fat tissue engineering goals in cellular agriculture. While spontaneously immortalized cells offer some advantages over genetically modified options, certain cell types are difficult to immortalize spontaneously, necessitating a genetic engineering approach. Gene editing technologies like CRISPR provide a targeted method to establish immortalized bovine satellite cells (BSCs) with precise control over genetic modifications, which may facilitate regulatory approval and consumer acceptance. The strategies used in gene editing to immortalize cells often modify genes like TP53^41–43^. TP53 (cellular tumor antigen), PTEN (phosphatase and tensin homolog) and SMAD4 (SMAD [Sma and Mad proteins] family member 4) regulate cell proliferation ^17,44,45^. In the present study, these genes were targeted for knock-out using CRISPR/Cas9 system to achieve immortalize BSCs^46,47^.

Two immortalized bovine satellite cell lines, CriBSC1 and CriBSC2, were successfully established, each demonstrating over 100 population doublings. In contrast, the two SMAD4 knockout cell lines and the PTEN knockout cell line failed to become immortalized. In the case of PTEN, as mentioned, this could be due to a failure of the mutations to knockout expression (as they did not reduce PTEN transcription), or in a functional downregulation at the protein level to fail to immortalize the cells. Future work should investigate this further. Compared to the CriBSC1 (TP53-gRNA2), which achieved nearly 150 total doublings, the TP53-gRNA1 cell line stopped proliferating after 91 total doublings. This suggests that the eighth exon of TP53’s DNA interaction sites is less critical for the immortalization of bovine satellite cells than the seventh exon of the DNA binding sites. Interestingly, CriBSC1 lost its differentiation capability, which may limit its application in cultivated meat production. However, it could still serve as a valuable model immortalized cell line for in-depth studies on bovine satellite cell differentiation. Importantly, CriBSC2 preserved both proliferation and differentiation during long-term culture, demonstrating growth and differentiation on both 2D and 3D environments. Future adaptation of these techniques for animal component-free scaffolds will be crucial for advancing cellular agriculture applications.

Both CriBSC1 and CriBSC2 underwent investigation using a range of biomarkers (PAX, MyoD, MyoG and MyHC) to elucidate cell proliferation and differentiation characteristics. In CriBSC1, the expression levels of MyoD and MyoG were relatively low during both proliferation and differentiation, contrasting with CriBSC2 where these markers were retained. Immunostaining further confirmed that CriBSC2 maintained differentiation capacity, while CriBSC1 completely lost this capacity. These findings suggest that simultaneous knockout TP53 and SMAD4 in BSCs leads to both immortalization and preservation of differentiation capacity. TP53 can inhibit myogenin transcription by directly binding to its promoter, thereby suppressing muscle cell differentiation^48^. SMAD4 is closely associated with restricting muscle differentiation in favor of proliferation, and its knockout has previously been shown to enhance terminal differentiation in satellite cells^49^. As such, it is possible that by knocking out TP53 and SMAD4 together, CriBSC2 are able to both maintain proliferation and differentiation capabilities. By knocking out the TP53 and SMAD4 together in CriBSC2, these pathways are preserved, maintaining both proliferation and differentiation capabilities. CriBSC2 also demonstrated the fastest doubling time, suggesting that downregulating TP53 and SMAD4 may synergistically promote cell proliferation through the cooperative action of cell cycle activators in these cells^46^.

The significant upregulation of pathways related to cell cycle and cell replication mechanisms in CriBSC2 compared to primary BSCs for all CriBSC2 passage timepoints in our pathway enrichment analysis results, suggests that targeting TP53 and SMAD4 may increase the proliferative abilities of BSCs. Additionally, the downregulation of early-stage MuSC activation and myogenic commitment and late-stage MuSC activation / myoblast differentiation markers (MYOD1, MYOG, TNNT2) and concurrent stability of biomarkers of quiescent or close-to-quiescent MuSCs (PAX3, SPRY1, HES1, GAS1, CALCR, KLF7), indicates that proliferating CriBSC2 maintain the features of MuSCs^50,51^. We propose to follow-up with additional experiments including whole genome sequencing analysis on this cell line over the course of several passages to assess the genetic stability of these cells. Further studies could also explore the genomic and transcriptomic differences in genetic stability between CriBSC2 and BSCs immortalized by different methods.

Compared to immortalized BSCs previously generated through constitutive overexpression of TERT and CDK4^10^, CriBSC2 presents potential regulatory and consumer benefits. CRISPR-mediated gene editing encounters fewer regulatory hurdles and enjoys improved consumer sentiment in many countries^11,12^. Even within stringent regulatory frameworks like the European Union, recent proposals indicate a pathway for regulatory approval of “new genomic techniques” such as CRISPR/Cas9 in food production, contrasting with the challenges faced by transgenic engineering methods used in our prior cell lines^10^. However, potential consumer acceptance for CriBSC2 for cultivated meat may face challenges due to the targeted genes being well-studied oncogenes, mutations of which are often linked with cancer *in vivo*^52^. Some media reports have already suggested concerns that the use of immortalized cells could imply consumption of cancerous tissues. This issue is primarily a communication challenge rather than a health risk, as noted by the Food and Agriculture Organization (FAO) of the United Nations, which concluded that consuming immortalized cells poses no credible pathway to harm^53^. From a scientific perspective, labeling immortalized cells as “cancerous” is inaccurate, as replicative immortality is just one of the hallmarks of cancer, which itself is a disease contextually dependent on a harmful *in vivo* conditions^54^. Nevertheless, addressing communication challenges and consumer concerns remains crucial. Future research should focus on consumer perception studies, direct comparisons of CriBSC2 cells with TERT/CDK4-expressing cells, transcriptomic comparison across different BSC types, and potential enhancements such as nutritional improvements and metabolic efficiency. Additionally, evaluating the stability, safety, and quality of CriBSC2 and the food derived from them will optimize their application in cultivated meat production. Further investigation into synergistic effects among the knockouts of PTEN, TP53 and SMAD4 in CriBSC2^55^, adaptation to serum-free media and culture on animal component-free scaffolds are critical steps towards real-world implementation for cultured meat^39,56^

In this study, CRISPR-mediated knockout of PTEN, TP53 and SMAD4 was explored to immortalize bovine satellite cells. CriBSC2, engineered with knockouts of PTEN, TP53 and SMAD4 together, exhibited robust proliferation and differentiation capabilities compared to the primary bovine satellite cells in long term passages. Transcriptomic analysis revealed significant changes in regulatory signaling pathways affected by these CRISPR knockouts, influencing downstream effectors and fundamental control processes. CriBSC1, with knockout of TP53 at the DNA binding site only, demonstrated proliferation capability, but lacked differentiation capability. Other cell lines with single knockout of PTEN, SMAD4 or TP53 at the DNA interaction sites failed to achieve immortalization, failing to pass the 100 total population doublings. Moreover, CriBSC2 showed promising growth capabilities when cultured on scaffolds. Overall, the genetically modified CriBSC2 cell line holds promise as a suitable cell candidate for cellular agriculture, pending thorough regulatory and consumer evaluations. The strategy employed in this study can also be adapted for immortalizing other types of animal cells for food-related applications.

## Supporting information

SI-Synergetic hallmark knockouts immortalize bovine muscle stem cells for cellular agriculture

## Data availability

Sequencing data are available at NCBI GEO. The GEO accession number is available upon request.

## Code availability

The full reproducible code of our RNA-Seq Analysis Pipeline is available upon request.

## Supplementary Information

Supplementary Table 1. Primer sequences for PCR clones and NGS analysis

Supplementary Table 2. Summary of clean reads and genes mapped to the reference genome from the primary BSCs and CriBSC2.

## Acknowledgements

Thanks to the USDA (2021-69012-35978), TUCCA consortium, New Harvest for supporting this work. The authors acknowledge the Tufts University High Performance Compute Cluster (https://it.tufts.edu/high-performance-computing) which was utilized for the research reported in this paper.

