## Supplementary material for "Synergetic hallmark knockouts immortalize bovine muscle stem cells for cellular agriculture": SI-Synergetic hallmark knockouts immortalize bovine muscle stem cells for cellular agriculture

Table 1 Primers used for the NGS testing.

| Name | Sequence (5' - 3') |
|---|---|
| PTEN-gRNA1-T-F | ATCACAGTTGCACAGTATCCC |
| PTEN-gRNA-T-R | GTCTCTGGTCCTTACTTCCCC |
| TP53-gRNA1-T-F | GGGTCCCCCTTTCTGATTGC |
| TP53-gRNA1-T-R | CTTCTTTGGCTGTGGAGAGGA |
| SMAD4-gRNA1-T-F | TGCTCCGGAAATTGGAGACAT |
| SMAD4-gRNA1-T-R | CTGTAGAAACCTGCCACTACA |
| TP53-gRNA2-T-F | AGATGGGTTATCGCCATTCCA |
| TP53-gRNA2-T-R | GGGAATGCAGAGAGAGGGAAC |
| SMAD4-gRNA2-T-F | GTCTCCTGTAGCTCCTGAGTATTGG |
| SMAD4-gRNA2-T-R | GCAGGGGTCAGCACATCTGG |

Table 2 Summary of clean reads and genes mapped to the reference genome from the primary BSCs and CriBSC2

| Sample name | Total reads | Mapping ratio (%) | Multi-mapped ratio (%) | Unique mapped ratio (%) | Unique mapped reads | Ratio (%) aligned to | | |
|---|---|---|---|---|---|---|---|---|
| | | | | | | coding (mRNA) regions | intergenic regions | intronic regions |
| BSCP4-A | 17,015,467 | 92.77 | 1.54 | 91.23 | 15,523,431 | 89.52 | 6.85 | 3.63 |
| BSCP4-B | 23,059,122 | 92.24 | 1.46 | 90.77 | 20,931,044 | 89.59 | 6.44 | 3.97 |
| BSCP4-C | 24,133,447 | 91.89 | 1.92 | 89.97 | 21,713,655 | 88.64 | 6.81 | 4.55 |
| CriBSC2-A | 27,866,750 | 92.17 | 1.95 | 90.52 | 25,139,391 | 89.01 | 6.43 | 4.56 |
| CriBSC2-B | 25,677,432 | 92.53 | 2.01 | 90.52 | 23,243,845 | 89.27 | 6.43 | 4.3 |
| CriBSC2-C | 24,538,849 | 92.66 | 1.94 | 90.72 | 22,261,042 | 89.61 | 6.34 | 4.04 |
